# Varying Effects of Common Tuberculosis Drugs on Enhancing Clofazimine Activity in vitro

**DOI:** 10.1101/112573

**Authors:** Shuo Zhang, Wanliang Shi, Jie Feng, Wenhong Zhang, Ying Zhang

**Affiliations:** Department of Molecular Microbiology and Immunology, Bloomberg School of Public Health, Johns Hopkins University, Baltimore, MD 21205, USA; Shandong Medicinal Biotechnology Centre, Shandong Academy of Medical Sciences, Jinan 250062, China; Department of Infectious Diseases, Huashan Hospital, Fudan University, Shanghai 200040, China

## Abstract

Clofazimine (CFZ), originally developed as an anti-tuberculosis (TB) drug in the 1950s 1, is commonly used to treat leprosy and also nontuberculous mycobacterial (NTM) infections. 2 Although CFZ has good activity against *Mycobacterium tuberculosis*, it was not used in the treatment of pulmonary TB mainly because it had the side effect of skin discoloration and there were other more effective drugs like isoniazid (INH), rifampin (RIF) and pyrazinamide (PZA) already available for the treatment of TB. 2 However, the increasing emergence of multi-drug-resistant TB (MDR-TB) has revived interest in the use of CFZ to treat MDR-TB. 2,3

Although resistance to CFZ has been shown to be mediated by mutations in *Rv0678,4,5 Rv1979c,* or *Rv2535c (PepQ)*,*5* the mode of action of CFZ has remained poorly understood. CFZ appears to have multiple effects on *M. tuberculosis* including interference with redox cycling,1 production of reactive oxygen species, and membrane destabilization or dysfunction. 6 7 CFZ is a bacteriostatic drug (MIC =0.06 μg/ml) with a slow action where it has little effect on the colony count until after 2 weeks.8 Heightened recent interest in this drug became apparent when CFZ added to the current MDR-TB regimen (called Bangladesh regimen), is associated with shortening of the lengthy treatment from 18-24 months to 9 months. 3 Moreover, CFZ was recently shown to shorten the treatment of drug susceptible TB from 6 months to 3 months when added to the standard TB treatment regimen consisting of INH, RIF, PZA, and ethambutol (EMB).9 These findings suggest that CFZ may have some unique activity on mycobacterial persisters and that certain TB drugs may synergize with CFZ or vice versa. However, so far, no information is available on the effect of other TB drugs on the activity of CFZ against *M. tuberculosis.* To address this question, in this study, we evaluated the effects of commonly used firstline and second line TB drugs on the activity of CFZ against stationary phase *M. tuberculosis* culture enriched in persisters in vitro in a drug exposure assay

A 3 week old stationary phase *M. tuberculosis* H37Rv culture (108–9 bacilli/ml) grown in 7H9 liquid medium containing 10% albumin-dextrose/catalase (ADC) was washed and diluted in 7H9 medium without ADC (5 x 10^6^ bacilli/ml), which was used for drug exposure studies with CFZ in combination with the commonly used firstline drugs (RIF, PZA, EMB) and important secondline drugs amikacin (AMK), moxifloxacin (MFX), levofloxacin (LEV), and para-amino salicylate (PAS). The drugs were dissolved in dimethyl sulfoxide (DMSO) or water as appropriate. INH was not included as it has no activity on the stationary phase *M. tuberculosis* culture. CFZ (1 μg/ml) was incubated in combination with the following drugs at their in vivo relevant achieveable blood concentrations: RIF (4 μg/ml), PZA (30 μg/ml at pH 6.0 or 6.8), EMB (3 μg/ml), amikacin (8 μg/ml), moxifloxacin (2 μg/ml), levofloxacin (8 μg/ml), and PAS (10 μg/ml) as CFZ containing two drug combinations, with single drug and drug-free controls, for various times (1, 4, 7, and 14 days) without shaking. After drug exposure, the surviving bacteria in the above treatment groups were washed to remove drug 85 s, diluted (undiluted, 1:10, and 1:100) and plated directly on drug-free 7H11 agar plates for colony forming unit (CFU) counts to assess the effect of drug exposure without subculture. After incubation at 37 °C for 4 weeks, the CFU values for different treatments were determined (see Table 1).

**Tabel 1.**
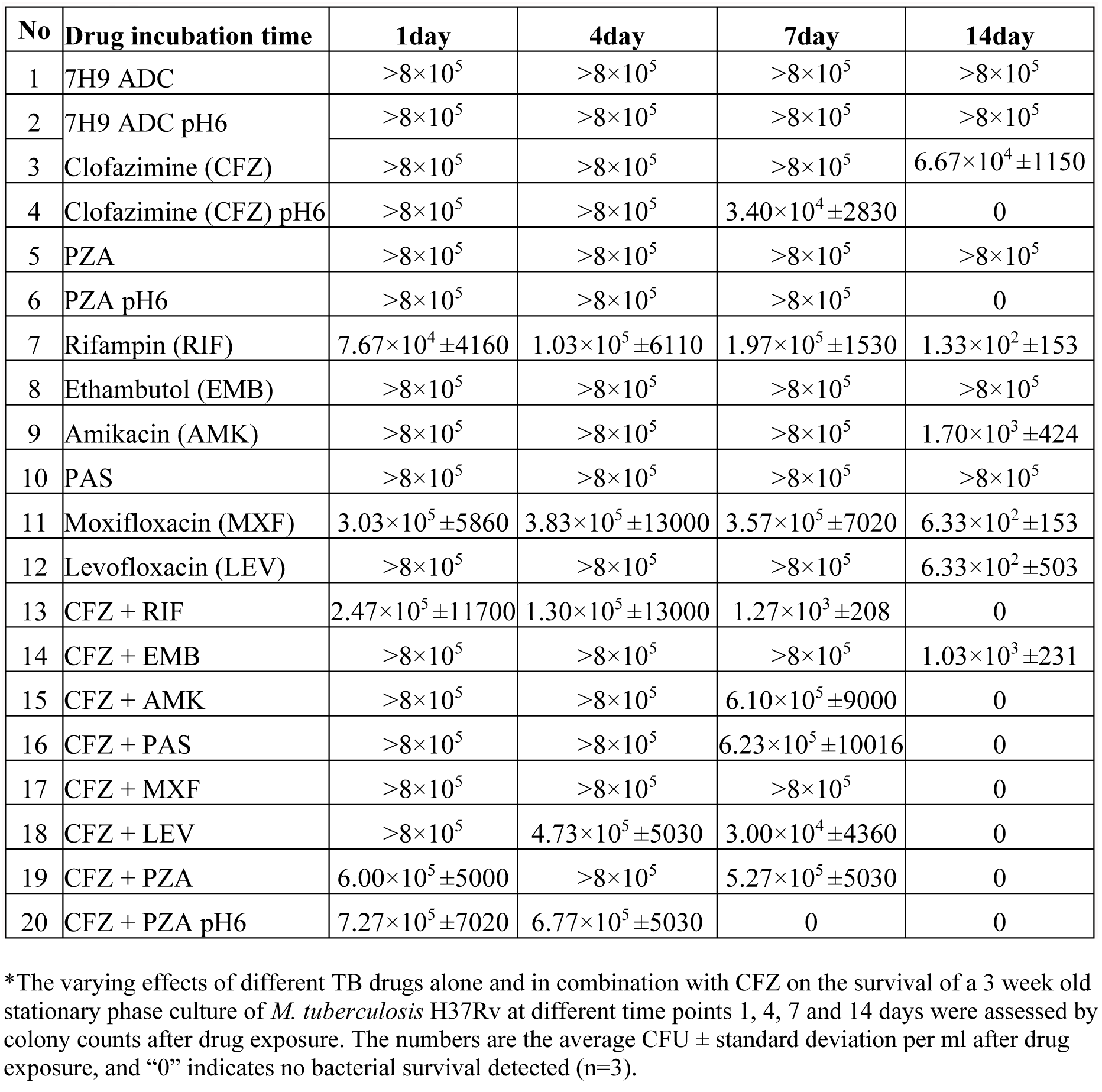
Varying effects of commonly used TB drugs on enhancing the activity of CFZ against stationary phase *M. tuberculosis* H37Rv *

It is of interest to note that the CFZ activity was significantly enhanced at acid pH6.0 as seen by less growth after 7 day drug exposure and no CFU remaining after 14 days (Table 1). In contrast, CFZ treatment alone at close to neutral pH 6.8 had poor activity against *M. tuberculosis* even after 14 day drug exposure (Table 1). The acid pH enhancement of CFZ activity was unexpected and not previously reported, and this is most likely caused by increased solubility of the poorly soluble CFZ (pKa=8.36) under acid pH. Future studies are needed to test this possibility in uptake experiments at acid pH with control drugs. Thus, it is possible that like PZA,10 the acid pH enhancement of CFZ activity may be relevant for in vivo situation during active inflammation that can produce acid pH. As a control, PZA at acid pH6.0 was more active than at close to neutral pH (6.8), as expected (Table 1). Except RIF which had some activity against the stationary phase culture, other single drugs (amikacin, moxifloxacin, levofloxacin, PAS, and CFZ and PZA at neutral pH) all had limited or poor activity against the 3 week old stationary phase culture (Table 1).

CFZ drug combination studies, we ranked the CFZ enhancement effects by commonly used firstline and secondline TB drugs. We found that PZA was by far the most active drug in enhancing the CFZ activity at acid pH6.0, followed by RIF, quinolones (moxifloxacin and levofloxacin), amikacin, and PAS in decreasing order of activity (Table 1). In contrast, cell wall inhibitor EMB had no apparent effect on enhancing CFZ activity (Table 1). Although we looked for other drugs that enhance CFZ activity, in fact, the combination effects can be said to be a reflection of mutual enhancements of CFZ and other TB drugs. Thus, it is noteworthy that we found in a separate study that CFZ could enhance PZA activity against *M. tuberculosis* (Niu H et al., submitted).

Despite the interesting observation of varying enhancement effects of CFZ activity exhibited by different TB drugs, the mechanisms involved remain to be determined and may differ in each specific case. For example, PZA enhancement of CFZ activity may be due to their concerted effect on disrupting the mycobacterial membranes which are a known persister target especially at acid pH.10 In addition, PZA may also enhance the CFZ activity through interfering with energy production via inhibition of PanD (aspartate decarboxylase) involved in CoA biosynthesis 11 such that it would deplete energy required to drive efflux of CFZ leading to increased accumulation of CFZ inside the cells to enhance its activity. We also found RIF increased the activity of CFZ (Table 1), and this could be due to the synergistic effect of RIF on causing inhibition of transcription of CFZ target leading to increased CFZ activity in the presence of RIF.

Gatifloxacin or moxifloxacin and CFZ a 129 re both included in the 9 month Bangladesh regimen for treating MDR-TB.3 It is of interest to note that we found quinolone drugs moxifloxacin and levofloxacin both enhanced the activity of CFZ against *M. tuberculosis* stationary phase cells (Table 1). In addition, we also observed amikacin enhanced the activity of CFZ. Our finding that amikacin enhanced the CFZ activity for *M. tuberculosis* is consistent with the previous finding that amikacin was shown to enhance CFZ activity against growing *M. abscessus* in vitro. 12 Our findings that multiple drugs including PZA, RIF (except cell wall inhibitor EMB) and secondline drugs (quinolones, amikacin and PAS) enhanced the activity of CFZ or vice versa, suggests a more general or broad effect of CFZ on *M. tuberculosis.* This observation is likely due to disruption of CFZ on bacterial memrbanes, 13 which is considered a good target for persister drugs. 14,15 In addition, our findings that many frontline and secondline drugs such as PZA, new generation quinolones (gatiflocaxin or moxifloxacin) and amikacin all enahnced the activity of CFZ also help to explain the high efficacy of the CFZ-containing 9 month Bangladesh regimen. 3

In summary, there is recent interest in understanding how CFZ might be involved in shortening the treatment of both MDR-TB and drug susceptible TB. The present study made a number of interesting observations that may help explain the unique ability of CFZ to shorten TB therapy, by demonstrating acid pH enhancement of CFZ activity, the varying degrees of enhancement of CFZ activity against statinary phase bacilli by different TB drugs, with PZA and RIF having the highest degree of enhancement, followed by quinolones (moxifloxacin and levofloxacin), amikacin and PAS. Future studies are needed to validate our in vitro fndings reported here in animal models of TB infection.

## ACKNOWLEDGEMENTS

The work was supported in part by NIH grants AI099512 and and AI108535.

